# Calcium signaling mediates mechanotransduction at the multicellular stage of *Dictyostelium discoideum*

**DOI:** 10.1101/2021.07.14.452436

**Authors:** Hidenori Hashimura, Yusuke V. Morimoto, Yusei Hirayama, Masahiro Ueda

## Abstract

Calcium acts as a second messenger and regulates cellular functions, including cell motility. In *Dictyostelium discoideum*, the cytosolic calcium level oscillates synchronously, and calcium signal waves propagate in the cell population during the early stages of development, including aggregation. At the unicellular phase, the calcium response through Piezo channels also functions in mechanosensing. However, calcium signaling dynamics during multicellular morphogenesis is still unclear. Here, live-imaging of cytosolic calcium levels revealed that calcium wave propagation, depending on cAMP relay, temporarily disappeared at the onset of multicellular body formation. Alternatively, the occasional burst of calcium signals and their propagation were observed in both anterior and posterior regions of migrating multicellular bodies. Calcium signaling in multicellular bodies occurred in response to mechanical stimulation. Both pathways, calcium release from the endoplasmic reticulum via IP3 receptor and calcium influx from outside the cell, were involved in calcium waves induced by mechanical stimuli. These show that calcium signaling works on mechanosensing in both the unicellular and multicellular phases of *Dictyostelium* using different molecular mechanisms during development.

**Summary statement:** Fluorescence imaging revealed that calcium signaling via both endoplasmic reticulum and extracellular pathways plays an important role in mechanosensing during the multicellular stage of *Dictyostelium*.

## Introduction

Ca^2+^ signals are essential for several types of biological activities (Clapham, 2007; Parekh, 2011). In multicellular organisms, the synchronized elevation of intracellular Ca^2+^ levels ([Ca^2+^]_i_) occurs in cell populations, and this “[Ca^2+^]_i_ burst” propagates as waves among cells (Berridge et al., 2003; Leybaert and Sanderson, 2012). This phenomenon has been reported in various cell types and biological activities, such as fertilization in eggs and wound repair of endothelial cells (Chifflet et al., 2012; Whitaker, 2006). [Ca^2+^]_i_ burst and wave propagation play key roles in orchestrating multiple cells *in vivo* and *in vitro* (Parekh, 2011). Cell-cell communication via Ca^2+^ signaling has been well investigated in animals, and it has revealed that Ca^2+^ waves are propagated by gap junction communication, or paracrine signaling (Leybaert and Sanderson, 2012). A factor evoking the [Ca^2+^]_i_ burst is a mechanical stimulus, and transduction of mechanical stimuli into Ca^2+^ signals is mediated by varied Ca^2+^ channels such as inositol trisphosphate (IP3) receptors, transient receptor potential (TRP) channels, and the stretch-activated ion channel Piezo (Canales et al., 2019; Coste et al., 2010; Fang et al., 2021; Prole and Taylor, 2019; Volkers et al., 2015; Yin and Kuebler, 2010). These channels are broadly conserved in eukaryotes including animals, plants, and amoebae (Coste et al., 2010; Volkers et al., 2015; Yin and Kuebler, 2010). Hence, there is a possibility that Ca^2+^ signaling is universally employed in various organisms for mechanosensing.

One example of Ca^2+^ wave propagation among populations of eukaryotic cells is cell-cell communication in the aggregation of social amoebae, *Dictyostelium discoideum*. Following starvation, *Dictyostelium* cells aggregate and form cell masses called mounds. Cells in a mound differentiate into either a prestalk or prespore cells and form migrating multicellular bodies called slugs. During aggregation, intercellular cAMP signaling, known as cAMP, relay organizes directed migration of cells (Gregor et al., 2010; Hashimura et al., 2019; Tomchik and Devreotes, 1981), and Ca^2+^ waves are propagated simultaneously among starved cells (Horikawa et al., 2010). It has been assumed that cAMP relay is essential for the coordination of collective cell migration through *Dictyostelium* development (Singer et al., 2019; Weijer, 1999); however, recent studies have shown that the dynamics of cAMP signaling shows transition after multicellular formation (Fujimori et al., 2019; Hashimura et al., 2019). In addition to Ca^2+^ wave propagation during aggregation (Horikawa et al., 2010), transient [Ca^2+^]_i_ elevation has been observed in mounds and slugs (Cubitt et al., 1995). These results suggest that [Ca^2+^]_i_ signaling such as synchronous [Ca^2+^]_i_ burst and wave propagation occurs not only during aggregation but also in the latter development stages of *Dictyostelium* cells, including mounds and slugs. However, the dynamics and molecular mechanisms of [Ca^2+^]_i_ signaling during the morphogenesis of *Dictyostelium* are still unclear.

In this study, the [Ca^2+^]_i_ signaling through the development of *Dictyostelium* cells was investigated. This approach revealed the transition of [Ca^2+^]_i_ signaling dynamics during the multicellular formation and the multicellular bodies of *Dictyostelium* have robust calcium signaling mechanisms in response to mechanical stimuli.

## Results

### Transition of calcium signaling dynamics in cell populations during the development of *Dictyostelium* cells

To investigate the relationship between the dynamics of calcium signals and multicellular formation in *Dictyostelium*, we monitored [Ca^2+^]_i_ dynamics during development with genetically-encoded calcium indicators (GECI). As previously reported (Horikawa et al., 2010), cells expressing the Förster resonance energy transfer (FRET) sensor YC-Nano15 (Kd = 15 nM) showed clear oscillations and wave propagation of fluorescence signals in aggregation streams (Fig. 1A, Fig. S1A, and Movie 1). Moreover, [Ca^2+^]_i_ dynamics was also investigated with the single-wavelength GECI, GCaMP6s (Chen et al., 2013; Pervin et al., 2018), to confirm whether the wave propagation of fluorescence signals of YC-Nano15 authentically reflected the [Ca^2+^]_i_ dynamics during development using the other GECI and avoiding the phototoxicity caused by exposure of violet-blue light excitation for YC-Nano15. In starved *Dictyostelium* cells, [Ca^2+^]_i_ transiently increases in response to external cAMP (Yumura et al., 1996) and the calcium channel IplA, which is the homologue of IP3 receptor, is essential for its elevation (Traynor et al., 2000). When chemotactic-competent cells expressing GCaMP6s were stimulated by cAMP, wild-type cells showed transient rapid elevation of fluorescence signals with a peak at 16 s after stimulation; however, cells lacking *iplA* show no increase of signals after stimuli (Fig S2A). Thus, GCaMP6s (Kd = 144 nM) (Chen et al., 2013) is functional in *Dictyostelium* cells and appropriate to visualize [Ca^2+^]_i_ dynamics during aggregation and sequential development. Oscillations of fluorescence signals and wave propagation were observed at the early aggregation and mound stages of cells expressing GCaMP6s (Fig. 1B–D, Fig. S1B–D, Movies 2–6). These signal propagation and oscillations were not observed in the populations of *iplA*^−^ cells during development (Fig. S2B, C, Movie 7), demonstrating that the periodic changes in GCaMP6s signals in developing *Dictyostelium* cells reflects [Ca^2+^]_i_ oscillations caused by cAMP signal relay. The period of oscillations at the early mound stage was significantly shorter than those at the early aggregation and late mound stages (*p* < 0.001) (Fig. 1E). The early and late mound stages observed in this study correspond to the loose and tight mound stages, respectively. The periods of [Ca^2+^]_i_ oscillations at the early aggregation, early and late mound stages were 5.29 ± 0.59, 2.95 ± 0.61, and 4.60 ± 0.89 min, respectively. These periods are consistent with those of [cAMP]_i_ oscillations (Hashimura et al., 2019). Wave propagation was observed until the late mound stage; however, signal oscillations and propagation in cell populations disappeared when the late mound began elongation, which is the onset of multicellular slug formation (Fig. 1 F–H, Movie 8). These indicate that [Ca^2+^]_i_ signal dynamics show transition during multicellular morphogenesis as well as cAMP signal dynamics (Hashimura et al., 2019).

**Fig. 1.**
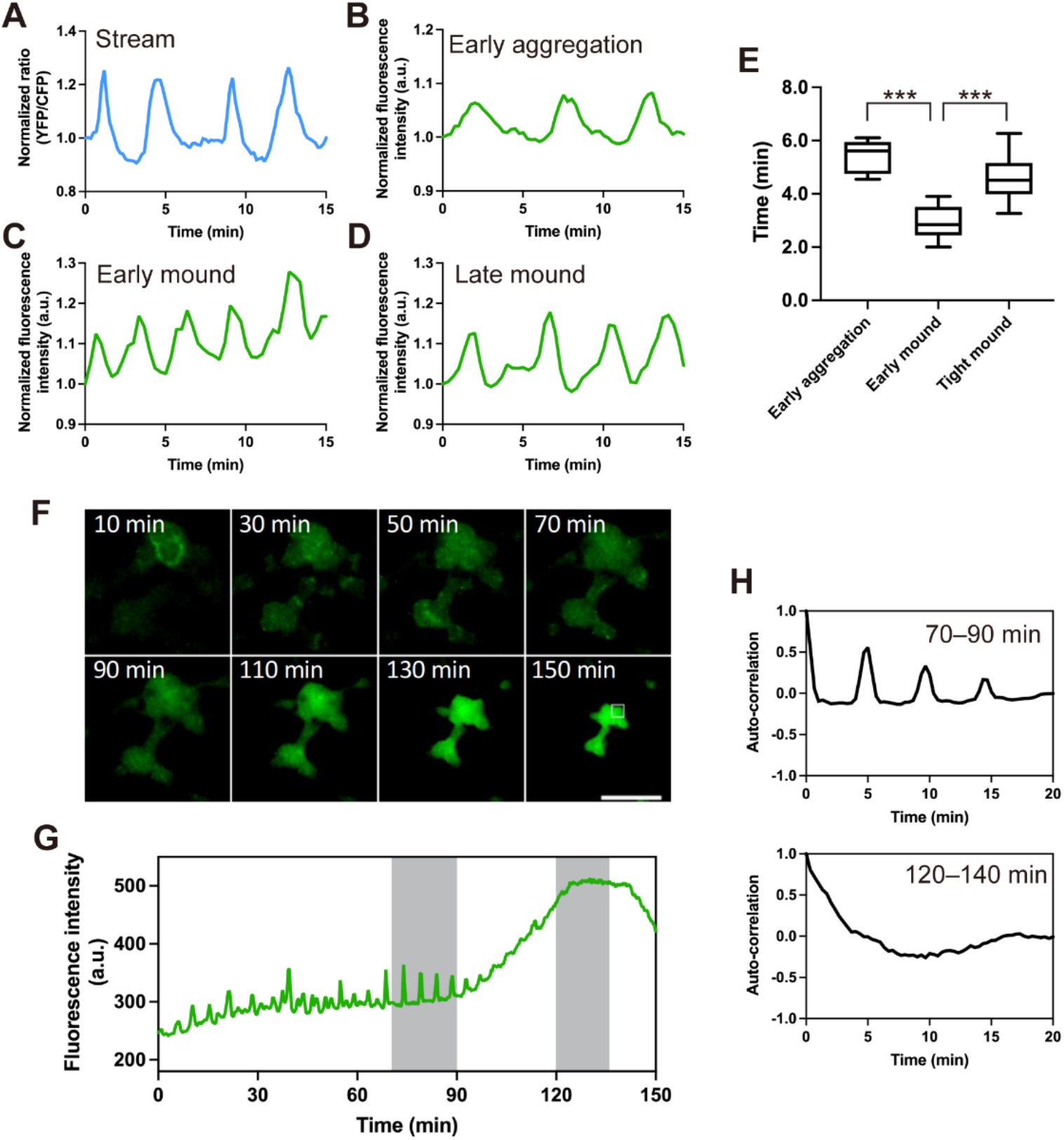
[Ca^2+^]_i_ signal dynamics at each developmental stage of *Dictyostelium* cells. Time course plots of (A) Förster resonance energy transfer (FRET) signals of YC-Nano15 or (B–D) mean fluorescence intensity of GCaMP6s in a region indicated by a white box (ROI) in Fig. S1. Size of ROI: A, 25 μm. B, 250 μm. C and D, 100 μm. (A) An aggregating stream. (B) Early aggregation. (C) An early mound. (D) A late mound. (E) Boxplot of periods of [Ca^2+^]_i_ oscillations at three developmental stages. The lower and upper error lines indicate the smallest and largest values, respectively. At each dataset, n = 13. ***; *p* < 0.001 (Wilcoxon rank sum test). (F) Fluorescence images of *Dictyostelium* cells expressing GCaMP6s during morphogenesis from the late mound to the finger stage. Scale bar, 500 μm. (G) Time course plot of mean fluorescence intensity of GCaMP6s in a 100 μm^2^ region indicated by a white box in F. (H) Autocorrelation of GCaMP6s signals at each development stage shown by gray bars in G. Note that the elevation of fluorescence intensity in the entire mound (90–150 min in F and G) is primarily due to the increase in thickness of the tissue rather than [Ca^2+^]_i_ elevation.

### Transient [Ca^2+^]_i_ burst and its propagation in migrating slugs

In the late stage of *Dictyostelium* development, the late mound elongates into a cylindrical structure called a finger, and finger subsequently falls over and starts to migrate as a slug. When monitoring [Ca^2+^]_i_ dynamics in migrating slugs using YC-Nano15, transient and rapid elevation of [Ca^2+^]_i_, namely “[Ca^2+^]_i_ burst” and its propagation was observed (Fig. 2A, B, Movie 9), although no wave propagation was observed at the finger stage (Fig. 1). Monitoring the signal using GCaMP6s also detected such transient signal propagation in migrating slugs, and the [Ca^2+^]_i_ bursts were observed in both the anterior and posterior part of slugs which can be regarded as the prestalk and prespore region, respectively (Fig. 2C–F, Movies 10, 11). When the [Ca^2+^]_i_ burst occurred in the slug, the velocity of slug migration transiently increased with a peak delay of approximately 2 min (Fig. 2B, Movie 9). The periodicity of [Ca^2+^]_i_ signals as observed in cell populations during aggregation and mound stages (Fig. 1) was not observed in migrating slugs, and slugs occasionally showed irregular [Ca^2+^]_i_ burst (Fig. 2 and S3). Thus, although periodic [Ca^2+^]_i_ signal propagations disappeared during the process of multicellular formation, the ability of Ca^2+^ signaling was maintained after slug formation, and the occasional propagation of [Ca^2+^]_i_ waves in migrating slugs affected the cooperative movement of cells.

**Fig. 2.**
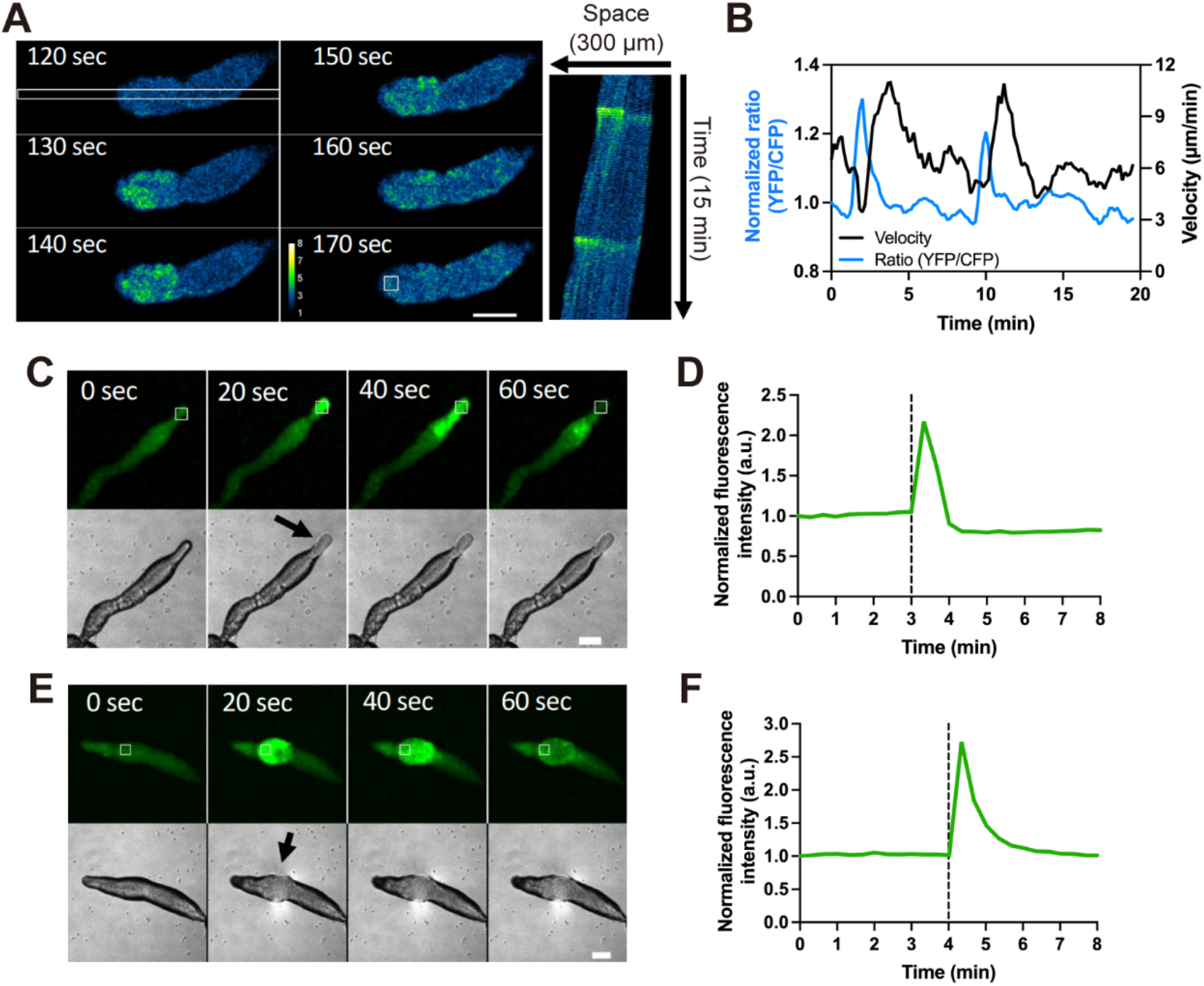
Intracellular Ca^2+^ levels ([Ca^2+^]_i_) burst in *Dictyostelium* slugs during migration. (A) Ratiometric images (YFP/CFP) of a slug expressing YC-Nano15 (Left). [Ca^2+^]_i_ burst at a tip of the slug expressing YC-Nano15 and its propagation toward the posterior region of the slug are shown. Scale bar, 50 μm. (Right) The kymograph of [Ca ^2+^]_i_ wave propagation in the region indicated by a white rectangle (10×300μm) in the left panel for 15 min duration. (B) Time-course plot of Förster resonance energy transfer (FRET) signals in the tip of the slug (blue line) and slug velocity (black line). The FRET signals of YC-Nano15 in a 15 μm^2^ region in the slug (white box in A) was measured. The curves of FRET signals and slug velocity were smoothed by a running average over four data points. (C and E) [Ca^2+^]_i_ burst at a tip (C) or posterior region (E) of the slug expressing GCaMP6s. Fluorescence images of GCaMP6s (upper panels)and differential interference contrast (DIC) images (lower panels) are shown. Scale bar, 100 μm. An arrow shows that the slugs are in contact with the agar surface. (D and F) Time course plot of the mean fluorescence intensity of GCaMP6s in a 50 μm^2^ region indicated by a white box in C and E. Black dashed lines indicate the time point when the slug was in contact with the agar surface.

### [Ca^2+^]_i_ burst and wave propagation in slugs are induced by mechanical stimulation

On closer observation, [Ca^2+^]_i_ burst in slugs occurred when a part of the slug touched the surface of the agar (Fig. 2C, E and Movies 10, 11). This proposes the possibility that rapid [Ca^2+^]_i_ elevation in slugs was induced in response to mechanical stimuli. To confirm this, [Ca^2+^]_i_ dynamics was monitored using GCaMP6s when the slug was subjected to mechanical stimulation. The slug developed on the agar was cut out with the agar, turned over onto the glass, sandwiched between the glass and the agar, and pressed from above with a 5 mm diameter plastic rod such that the slug was not crushed, and the entire slug was stimulated evenly (Fig. S4A). In all tests using wild-type expressing GCaMP6s, [Ca^2+^]_i_ in the slug increased transiently with a peak at 25.0 ± 4.1 s after all stimulation (n = 9) (Fig. 3A, B and Movie 12). Additionally, when the tip of a slug was pricked using a micropipette (Fig. S4B), [Ca^2+^]_i_ burst was induced and signal waves were propagated toward the posterior region (Fig. 3C, D and Movie 13). A similar response was observed when the posterior region of the slug was stimulated (Fig. 3E, F and Movie 14). These indicate that [Ca^2+^]_i_ burst and wave propagation in the slug occurs in response to mechanical stimulation. In addition, this mechanical response can be applied to either the anterior or posterior regions of the slug.

**Fig. 3.**
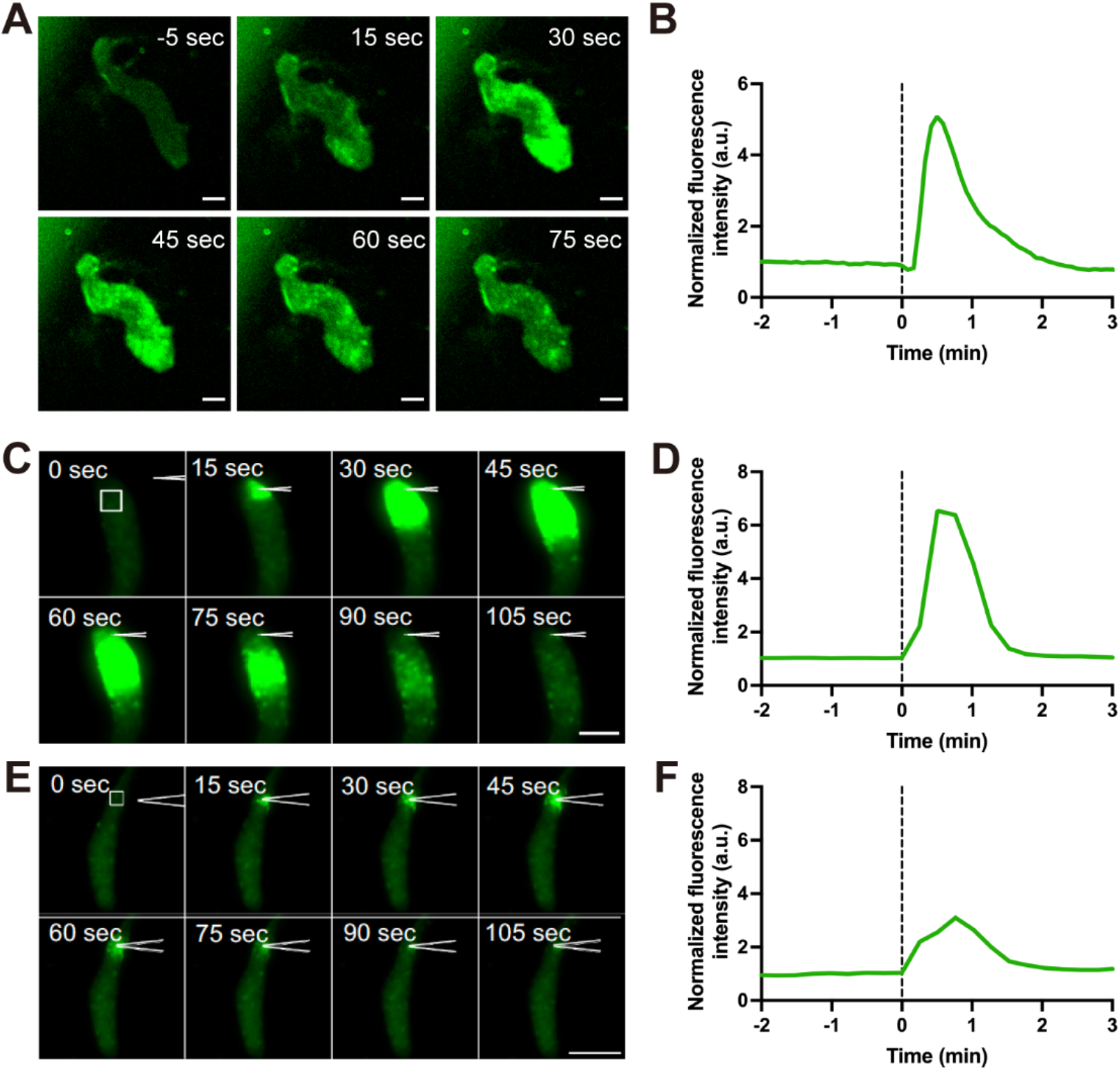
Intracellular Ca^2+^ levels ([Ca^2+^]_i_) burst in *Dictyostelium* slugs induced by mechanical stimulation. (A) Representative fluorescence images of a slug expressing GCaMP6s mechanically stimulated with a plastic rod. Scale bar, 50 μm. (B) Time course plot of the mean fluorescence intensity of GCaMP6s in a 25 μm^2^ ROI of the anterior region in A. (C and E) [Ca^2+^]_i_ burst at the tip (C) or posterior region (E) of the slug expressing Dd-GCaMP6s after mechanical stimulation by pricking with a micropipette. Fluorescence images of slugs expressing Dd-GCaMP6s are shown. Anterior part of the slug faces the top (C) or bottom (E) side of images. Scale bar, (C) 50 and (E) 100 μm. (D and F) Time course plot of the mean fluorescence intensity of GCaMP6s in a 25 μm^2^ region indicated by a white box in C and E, respectively. Black dashed lines indicate the time point of mechanical stimulation.

### IplA Ca^2+^ channel is involved in calcium signaling in response to mechanical stimulation in the slug

At the unicellular phase of *Dictyostelium*, the IP3 receptor IplA, which is localized in the endoplasmic reticulum (ER), is involved in calcium signaling, and responsible for [Ca^2+^]_i_ elevation in response to mechanical stimuli (Lombardi et al., 2008). To confirm whether IplA is involved in [Ca^2+^]_i_ burst induced by mechanical stimulation in a slug, [Ca^2+^]_i_ signal responses to mechanical stimuli in slugs lacking *iplA* was investigated. When *iplA*^−^ slugs were stimulated with a plastic rod, [Ca^2+^]_i_ bursts occurred; however, the percentage of slug that responded dropped to 46% (n = 39), and the response peaked at 15.7 ± 7.9 s (n = 18), earlier than in the wild type (Fig. 3, 4). Calcium response was also observed when *iplA*^−^ slugs bumped into the agar as well as the wild type (Fig. S5). These results suggest that [Ca^2+^]_i_ burst and wave propagation in response to mechanical stimuli are partially mediated by the IplA channel.

**Fig. 4.**
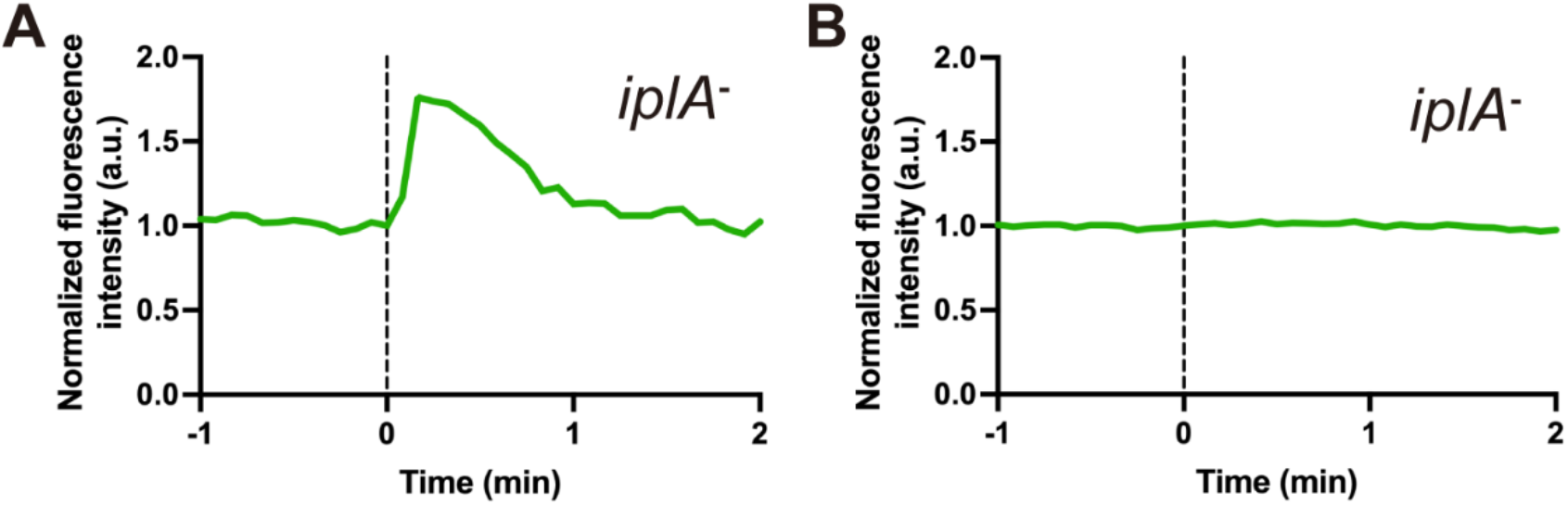
Intracellular Ca^2+^ levels ([Ca^2+^]_i_) burst in *iplA*^−^ slugs induced by mechanical stimulation. [Ca^2+^]_i_ was monitored in the slug of IplA null cells expressing GCaMP6s after mechanical stimulation with a plastic rod. (A) Representative time course plot of the mean fluorescence intensity of GCaMP6s in a slug showing [Ca^2+^]_i_ burst. (B) Representative time course plot of the mean fluorescence intensity of GCaMP6s in a slug showing no [Ca^2+^]_i_ response. Black dashed lines indicate the time point of mechanical stimulation.

### Calcium influx from outside the cell allows for a rapid response to mechanical stimuli

Deletion of IplA did not completely abolish the calcium response of the slug to mechanical stimulation (Fig. 4), indicating that another calcium pathway contributes to mechanosensing. It has been reported that extracellular Ca^2+^ influx via the Piezo channel homolog is important for mechanosensing at the unicellular stage of *Dictyostelium* (Srivastava et al., 2020). To investigate whether Ca^2+^ influx from outside the cell occurs even in the multicellular stage, calcium response was monitored with the agar medium containing ethylene glycol-bis(β-aminoethyl ether)-N,N,N’,N’-tetraacetic acid (EGTA), a Ca^2+^ chelating agent. All multicellular bodies overlayed with agar containing 1 mM EGTA showed an increase in [Ca^2+^]_i_ in response to mechanical stimuli (n = 13) (Fig. 5A, B). However, in the presence of EGTA, the response peak was significantly slower at 67.7 ± 16.8 seconds (*p* < 0.001) (Fig. 5B, E). Additionally, *iplA*^−^ slugs in the presence of 1 mM EGTA did not show any calcium response to mechanical stimulation (n = 13) (Fig. 5C). These and *iplA*^−^ deficient results indicate that Ca^2+^ influx from extracellular sources allows a fast response, whereas IplA is essential for the response from intracellular sources (Fig. 4, 5C). We constructed the PzoA null strain and confirmed that PzoA is essential for Ca^2+^ influx from extracellular sources by mechanical stimulation in unicellular cells as previously reported (Fig. S6) (Srivastava et al., 2020). In contrast, multicellular bodies of *pzoA*^−^ strain responded similarly to wild-type cells, suggesting that Ca^2+^ influx occurs from other pathways during the multicellular phase (Fig. 5D, E).

**Fig. 5.**
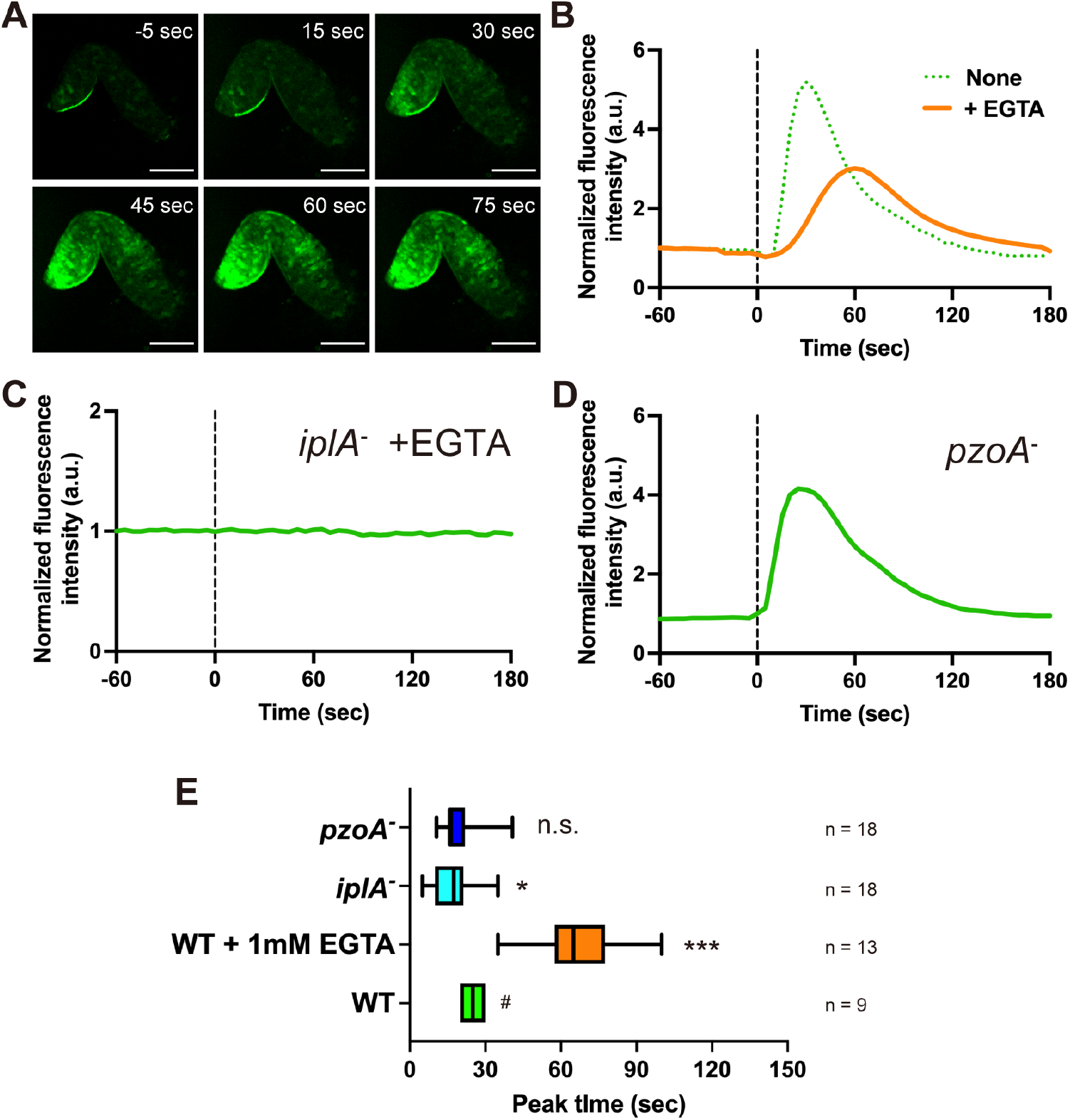
Effect of extracellular Ca^2+^ on intracellular Ca^2+^ levels ([Ca^2+^]_i_) burst in *Dictyostelium* slugs induced by mechanical stimulation. (A) A wild-type slug was covered with a piece of agar containing 1 mM aminoethyl ether)-N,N,N’, N’-tetraacetic acid (EGTA). Representative fluorescence images of GCaMP6s in a slug mechanically stimulated with a rod on EGTA agar. Scale bar, 100 μm. (B) Orange line shows time course plot of the mean fluorescence intensity of GCaMP6s in a 25 μm^2^ ROI of the anterior region in A. The green dashed line shows the plot of fluorescence intensity without EGTA in Fig. 3B. (C) Time course plot of the mean fluorescence intensity of GCaMP6s in the slug of IplA null cells with agar containing 1 mM EGTA. (D) Time course plot of the mean fluorescence intensity of GCaMP6s in the slug of PzoA null cells without 1 mM EGTA. (E) Box plots of the peak time of [Ca^2+^]_i_ burst after mechanical stimuli. Lower and upper error lines indicate the smallest and largest values, respectively. Average values were compared with the peak time of the wild-type slug without EGTA (#). *, *p* < 0.05; ***; *p* < 0.001 (Two tailed *t*-test). n.s. indicates no significant difference.

## Discussion

In *Dictyostelium* cells, transient [Ca^2+^]_i_ changes have been observed in the mound and slug stages (Cubitt et al., 1995); however, the actual dynamics of Ca^2+^ signaling has not been clarified, because most previous studies of calcium signaling have focused on the stages up to cell aggregation (Horikawa et al., 2010; Nebl and Fisher, 1997; Schlatterer et al., 1992; Traynor et al., 2000). In this study, live imaging of [Ca^2+^]_i_ during the development of *Dictyostelium* with highly-sensitive GECIs revealed that synchronized [Ca^2+^]_i_ elevation and its propagation in cell populations occur continuously at the aggregation and mound stages; however, they temporarily disappear during multicellular formation. Ca^2+^ wave propagation depends on cAMP relay at the early aggregation and mound stages, and cAMP wave propagation disappears during tip elongation of the late mound (Fujimori et al., 2019; Hashimura et al., 2019). The cAMP signal has been shown to induce transient [Ca^2+^]_i_ elevation (Abe et al., 1988; Nebl and Fisher, 1997). Therefore, changes in the dynamics of [Ca^2+^]_i_ during development follow the transition of cAMP relay. Alternatively, we found that [Ca^2+^]_i_ burst and its propagation occasionally occurred in slugs. When [Ca^2+^]_i_ waves were propagated in the migrating slug, velocity of the slug transiently increased. Ca^2+^ signaling affects both cell movement at the unicellular stage and slug behavior (Dohrmann et al., 1984; Fache et al., 2005; Lombardi et al., 2008). Thus, [Ca^2+^]_i_ wave propagation is involved in coordinated movements of multiple cells throughout development. Calcium wave propagation and its effect on cell behavior are also well known in animal cells, and gap-junction is essential for cell-cell signaling (Leybaert and Sanderson, 2012). Given that *D. discoideum* has no homologue of gap junction components (Johnson et al., 1977; Kaufmann et al., 2012), the mechanism for calcium signal propagation in multicellular bodies is gap junction independent. Another possible mechanism of calcium wave propagation is ATP-mediated paracrine signaling (Leybaert and Sanderson, 2012). It has been reported that *Dictyostelium* cells release ATP as an extracellular signal (Sivaramakrishnan and Fountain, 2015). Extracellular ATP causes an increase in [Ca^2+^]_i_ in *Dictyostelium* cells via P2X receptors and polycystin-type Trp channels that are either the ATP receptor or closely coupled to ATP (Ludlow et al., 2008; Traynor and Kay, 2017). Recently, it has been suggested that ATP levels contribute to differentiation during the multicellular phase (Hiraoka et al., 2020). This implies that slug cells secrete ATP, which might trigger wave propagation of elevated [Ca^2+^]_i_ in tissue.

*Dictyostelium* cells show [Ca^2+^]_i_ elevation in response to cAMP signals and mechanical stimuli at the unicellular phase (Artemenko et al., 2016; Lombardi et al., 2008; Srivastava et al., 2020). Our assay showed that [Ca^2+^]_i_ burst and its propagation were induced by mechanical stimuli at the slug stage. The IplA Ca^2+^ channel is involved in [Ca^2+^]_i_ elevation in response to mechanical stimuli at the unicellular phase (Artemenko et al., 2016; Lombardi et al., 2008; Srivastava et al., 2020). In slugs of the *iplA*^−^ strain, the response efficiency of calcium signaling to mechanical stimulation was reduced, suggesting that IplA is responsible for increasing the certainty of response to mechanical stimuli. The IplA channel is essential for Ca^2+^ dependent flow-directed motility; however, not for chemotactic migration toward cAMP gradients (Lusche et al., 2012). This suggests that the cAMP signaling pathway and the IplA mediated Ca^2+^ signaling pathway affect the downstream independently each other. This proposition is supported by the observation that there is no clear defect in the development of *iplA*^−^ cells under laboratory conditions (Movie 7). However, mechanosensing may play important roles in efficient morphogenesis in natural environments where cells of soil-living amoebae *Dictyostelium* are exposed to various stimuli and physical barriers (Bonner and Lamont, 2005). Therefore, mechanical stimulation response of Ca^2+^ signaling via IplA would be important in natural environments (Movie 2). Alternatively, the calcium signaling response to mechanical stimulation was not completely abolished in *iplA*^−^ slugs, suggesting that other signal pathways are involved in the elevation of cytosolic [Ca^2+^]_i_ in the mechanical response of slugs. In higher eukaryotes, the stretch-activated Ca^2+^ permeable ion channel Piezo is involved in mechanical stimulus responses (Coste et al., 2010; Fang et al., 2021). Recently, it has been reported that *D*. *discoideum* has the homologue of Piezo, PzoA, and disruption of the *pzoA* gene causes the defect of [Ca^2+^]_i_ response to mechanical stimuli at the unicellular phase (Srivastava et al., 2020). Additionally, mutant cells lacking PzoA can develop normally under laboratory conditions; however, a defect is observed in chemotactic migration under confined conditions (Srivastava et al., 2020). Slugs with cells lacking *pzoA* did not show a substantial difference in calcium response from the wild type. Notably, the Piezo channel, which works in higher multicellular organisms, works only in the unicellular phase in *Dictyostelium*. Even within *Dictyostelium*, the system changes differently, with Piezo acting as the main pathway during the unicellular phase, and pathways from the extracellular and ER during the multicellular phase. In IplA-null cells, the [Ca^2+^]_i_ response was faster than in the wild type. This indicates that the apparent single-peak [Ca^2+^]_i_ burst is a mixture of a fast extracellular Ca^2+^ influx and a slower yet more efficient response from the ER. The delay in the calcium response from the ER compared to extracellular signals may be due to the fact that the signal from mechanoreceptors in the plasma membrane is transmitted via IP3 signaling. Moreover, in the unicellular phase of IplA-null cells, no calcium response to mechanical stimuli was detected, even though IplA is not involved in blebbing, which is regulated by calcium signaling related to mechanical stimulation (Srivastava et al., 2020). In human colorectal carcinoma cell line (DLD1 cells), membrane blebbing is regulated by store-operated calcium entry, which is controlled by ER proteins (Aoki et al., 2021). Alternatively, although no homologue of STIM has been found in *Dictyostelium* (Prakriya and Lewis, 2015), unknown store-operated calcium channels (SOCs) may be transducing mechanical stimuli.

[Ca^2+^]_i_ burst at both the tip and posterior regions in slugs indicates the ability of rapid [Ca^2+^]_i_ elevation in both prestalk and prespore cells. Previous studies has shown that [Ca^2+^]_i_ in the anterior part of the slugs is higher than that in the posterior part (Cubitt et al., 1995; Yumura et al., 1996). However, in our study, such a difference was not observed in slugs at the stationary phase that did not show any [Ca^2+^]_i_ burst. As slugs migrate with their tips protruding up and down (Breen et al., 1987), the frequency of [Ca^2+^]_i_ bursts evoked by mechanical stimuli is higher in the anterior region that that of the posterior. Thus, it may have been frequently observed in previous studies that [Ca^2+^]_i_ is higher in the anterior part of the slug than in the posterior part.

In conclusion, we revealed that [Ca^2+^]_i_ burst and its propagation in populations of *Dictyostelium* cells occur dependently on cell-cell communication via diffusible chemical signals during the early developmental stage. Following multicellular formations, such Ca^2+^ signaling is triggered by mechanical stimuli. The system of Ca^2+^ signaling in response to mechanical stimuli is conserved broadly in higher eukaryotes, animals, and prokaryotes such as *Escherichia coli* (Bruni et al., 2017; Leybaert and Sanderson, 2012; Wakai et al., 2021). We observed that social amoebae *D*. *discoideum* belonging to Amoebozoa uses Ca^2+^ signaling as a mechanosensing signal at the multicellular phase similarly to the unicellular phase (Srivastava et al., 2020); however, the molecular mechanism is different. Thus, this study demonstrates that mechanochemical signal transduction via Ca^2+^ signaling is a universal system for response to mechanical stimuli and can be applied in any cell type or state. In this study, the extracellular Ca^2+^ pathway associated with mechanical stimulation in the multicellular bodies of *D*. *discoideum* has not been identified. This is because even though Ca^2+^ act as a signal across species, the molecular mechanisms differ between species. Further studies are required to clarify conserved and specific molecular mechanisms, respectively.

## Material and Methods

### Cell strains and culture conditions

*Dictyostelium discoideum* strains used in this study are listed in Table S1. The *pzoA*^−^ strain was constructed using the vector pKOSG-IBA-dicty1 (iba) following the manufacturer’s instructions (Wiegand et al., 2011). The 5’ region and 3’ region of flanking sequences were generated via polymerase chain reaction (PCR) and cloned into pKOSG-IBA-dicty1 (Fig. S6A). Primer pairs used for PCR were pzA_KO_LA1/pzA_KO_LA2 (5’) and pzA_KO_RA1/pzA_KO_RA2 (3’) (Table. S2). *pzoA* gene disruption was confirmed via PCR. Cells were grown axenically in HL5 medium (Formedium, UK) in culture dishes at 21 °C. Transformants were maintained at 20 μg mL^−1^ G418 (Fujifilm Wako, Japan) or 10 μg mL^−1^ BlasticidinS (Fujifilm Wako, Japan).

### Plasmid construction and genetic manipulation

Plasmids used in this study are listed in Table S3. pHK12neo_Dd-GCaMP6s was constructed by insertion of synthesized GCaMP6s fragments (GenScript) into the BglII and SpeI sites of pHK12neo. The codon usages of the GCaMP6s sequence were optimized to those of *D*. *discoideum* for efficient protein expression in *Dictyostelium* cells. The wild-type strain and mutant cells were transformed with ~1.5 μg plasmid via electroporation (Kuwayama et al., 2008), and transformants were selected with G418 or BlasticidinS.

### Instruments for image acquisition and analysis

In all experiments, cells were observed at 22 °C. Confocal images were taken using a confocal laser microscope (A1 confocal laser microscope system, Nikon, Japan) with an oil immersion lens (Plan Fluor 40×/1.30 NA, Nikon) or an inverted microscope (Eclipse Ti, Nikon) equipped with a CSU-W1 confocal scanner unit (Yokogawa), two sCMOS cameras (ORCA-Flash4.0v3, Hamamatsu Photonics, Japan) and an oil immersion lenses (Plan Apo 60×/1.40 NA or CFI Apo TIRF 60×/1.49, Nikon). GCaMP6s and YC-Nano15 were excited using a 488 and 440 nm solid-state CW laser, respectively. Epifluorescence imaging was taken using an inverted epifluorescence microscope (IX83, Olympus, Japan) equipped with a 130 W mercury lamp system (U-HGLGPS, Olympus), sCMOS cameras (Zyla4.2, Andor Technology or Prime 95B, Photometrics, USA) and objective lenses (UPLSAPO 4×/0.16 NA, UPLSAPO 10×/0.40 NA, UPLSAPO 20×/0.75 NA, Olympus). Cells expressing GCaMP6s were observed using fluorescence mirror units U-FGFP (Excitation BP 460–480, Emission BP 495–540, Olympus). All images were processed and analyzed using Fiji and R software. The period of an oscillation of GCaMP6s signals was calculated by averaging the difference between the peaks of the oscillation. Data with at least three peaks in the oscillation were used for the analysis. In general, fluorescence intensities of GCaMP6s and the ratio of YFP/CFP channels of YCNano15 were normalized with values at t = 0.

### Live image of [Ca^2+^]_i_ dynamics during *Dictyostelium* development

Image acquisition of *Dictyostelium* development was performed as reported by a previous study (Hashimura et al., 2019). To induce development upon starvation, cells at the exponential phase (1.5–3 × 10^6^ cells ml^−1^) were harvested and washed three times in KK2 phosphate buffer (20 mM KH_2_PO_4_/K_2_HPO_4_, pH 6.0). To observe all developmental stages, cells were plated on the entire surface of 2% water agar (2% w/v Difco Bacto-agar in ultrapure water) at a density of 5–7 × 10^5^ cells cm^−2^ on a 35-mm plastic dish (Iwaki, Japan) and incubated at 21 °C. Thereafter, plates were filled with liquid paraffin (Nacalai Tesque, Japan) to avoid light scattering and placed on the stage of the microscope for image acquisition. Additionally, the “2D slug” method (Bonner, 1998; Rieu, Barentin, Sawai, Maeda, & Sawada, 2004) was applied for observing slug migration without three-dimensional scroll movement (Fig. 4A and S2). One microliter cell suspension (4 × 10^7^ cells mL^−1^) was dropped on 2% water agar plates with 2 μL liquid paraffin. A coverslip was placed over the suspension and incubated at 21 °C for longer than 15 h.

### Validation of GCaMP6s as an indicator of [Ca^2+^]_i_ changes in chemotactic-competent *Dictyotelium* cells

*Dictyostelium* cells expressing Dd-GCaMP6s were suspended in 1 mL developmental buffer (DB: 5 mM Na/KPO_4_, 2 mM MgSO4, 0.2 mM CaCl2, pH 6.5) at a density of 5 × 10^5^ cells mL^−1^ and incubated for 1 h. Thereafter, cells were exposed to 100 nM cAMP pulses given at 6 min intervals for a subsequent 5 h. Following starvation with cAMP pulse treatment, cells were washed three times with 1 mL DB and resuspended in DB at a density of 10^6^ cells mL^−1^. Forty microliters of cell suspension was deposited onto a glass bottom dish. Cells were stimulated by adding 160 μL of 12.5 μM cAMP (Sigma Aldrich, USA) solution to the cell droplet (final concentration, 10 μM). During stimulation, fluorescent images of starved cells were taken using the confocal microscope at 5 s interval. Averaged fluorescence intensities of GCaMP6s in 5 μm^2^ regions positioned within the cytosol were estimated at each time point.

### Monitoring of [Ca ^2+^]_i_ response in slugs to mechanical stimulation

To observe the response of migrating slugs to mechanical stimulation, 5 μL of cell suspension at a density of 4 × 10^7^ cells mL^−1^ was deposited on 2% water agar with or without 1 mM EGTA and incubated at 21 °C for 12–15 h. Following slug formation, a piece of agar with slugs was cut out and placed upside down on a spacer attached to a 35 mm glass bottom dish (12 mm diameter glass, Iwaki). The spacer was filled with liquid paraffin to prevent desiccation during observation and avoid light scattering. A slug covered by agar was pushed with a 5 mm diameter plastic rod using a micromanipulator system (MM-94 and MMO-4, Narishige, Japan) (Fig. S4A). In the micropipette assay, a piece of agar with slugs was cut out and placed directly on a 35 mm glass bottom dish (12 mm diameter glass, Iwaki). A wet paper was placed in the dish and the agar piece was covered with liquid paraffin. A Femtotip microcapillary (1 μm tip diameter, Eppendorf, Germany) was mounted onto a Femtojet pump and micromanipulator (Eppendorf), and the pipette was pricked on the slug by manual operation with the manipulator (Fig. S4B). During mechanical stimulation, fluorescence images of slugs expressing GCaMP6s were acquired at 5 s interval using the epifluorescence microscope.

## Acknowledgements

We acknowledge Yuko Baba for technical assistance and Takuo Yasunaga for continuous support and encouragement. H.H. was supported by the RIKEN JRA program.

## Author contributions

H.H. conceived and designed the study, performed the experiments, analyzed the data, and wrote the manuscript. Y.H. performed the experiments and analyzed the data. M.U. designed the study, contributed to the interpretation of the data analysis. Y.V.M. designed the study, performed the experiments, analyzed the data and wrote the manuscript.

